# *srahunter*: a user-friendly tool to speed up and simplify data downloading from NCBI SRA

**DOI:** 10.1101/2024.03.19.585745

**Authors:** Enrico Bortoletto, Riccardo Frizzo, Paola Venier, Umberto Rosani

**Affiliations:** Department of Biology, University of Padova, 35121 Padova, Italy

## Abstract

Easy access and use of vast datasets are paramount for advancing scientific discovery in steadily expanding studies based on high-throughput sequencing (HTS). The Sequence Read Archive (SRA) is a publicly accessible repository currently holding a huge amount of HTS reads, as part of the International Nucleotide Sequence Database Collaboration (INSDC). However, accessing, downloading, and managing data and metadata efficiently can be challenging. Here, we introduce *srahunter*, a tool designed to simplify data and metadata acquisition from SRA. Developed with Python, *srahunter* leverages the core functionalities of *SRA Toolkit* and *Entrez Direct*, to enable automated downloading, smart data management, and user-friendly metadata integration through an interactive HTML table https://github.com/GitEnricoNeko/srahunter. Compared to existing tools, *srahunter* increases the efficiency of metadata retrieval by reducing the technical barriers to SRA data and streamlining the handling of SRA datasets, and can therefore accelerate the development of genomics and multiple ‘omics research.

## 1. Introduction

In the fast-evolving field of high-throughput sequencing (HTS), the availability of high-quality data is crucial for testing scientific hypotheses and driving new findings. As sequencing technologies advance, the resulting data increases exponentially, enabling researchers to explore complex biological questions. The Sequence Read Archive (SRA) established by the National Center for Biotechnology Information (NCBI) is the largest public repository of high-throughput sequencing data [1–3]. In 2023 the SRA database hosted more than 25.6 petabases from 14.8 million publicly available samples, indicating the rapid and exponential growth of HTS-based research encompassing the different fields of biology [3,4]. However, efficient access, download, and management of HTS data and metadata poses significant challenges, primarily due to the abundance of archived data and, secondly, to the complex structure of the sequencing data and associated metadata [1].

NCBI *Entrez direc*t [5] and *Sra Toolkit* (https://github.com/ncbi/sra-tools) are command-line tools developed by the NCBI SRA staff that allow the download of data and metadata from SRA. Both tools offer a set of options that may require users to spend time familiarising with them. Indeed, setting all parameters and identifying the best way to process the data requires time and a medium level of coding capabilities, especially for an extensive list of accessions.

Recognizing the possible bottlenecks, we have designed a tool to simplify data and metadata acquisition processes from the SRA, named *srahunter*. Based on *Sra toolkit* and *Entrez direct, srahunter* enables researchers to streamline data retrieval and handling via automated download, intelligent data management, and user-friendly metadata integration, including an interactive HTML table as standard output.

## 2. Functionality and Usage

*srahunter* takes as input an SRA accession list and successfully downloads datasets produced by different sequencing platforms, including the widely used short read platforms, like Illumina and DNBSEQ as well as long read sequencers, like Pacific Biosciences (PacBio) and Oxford Nanopore Technology (ONT). *srahunter* is composed of three different modules, named download, metadata, and fullmetadata (Figure 1). All three modules take the SRA Accession list of datasets to be retrieved as standard input.

**Figure 1.**
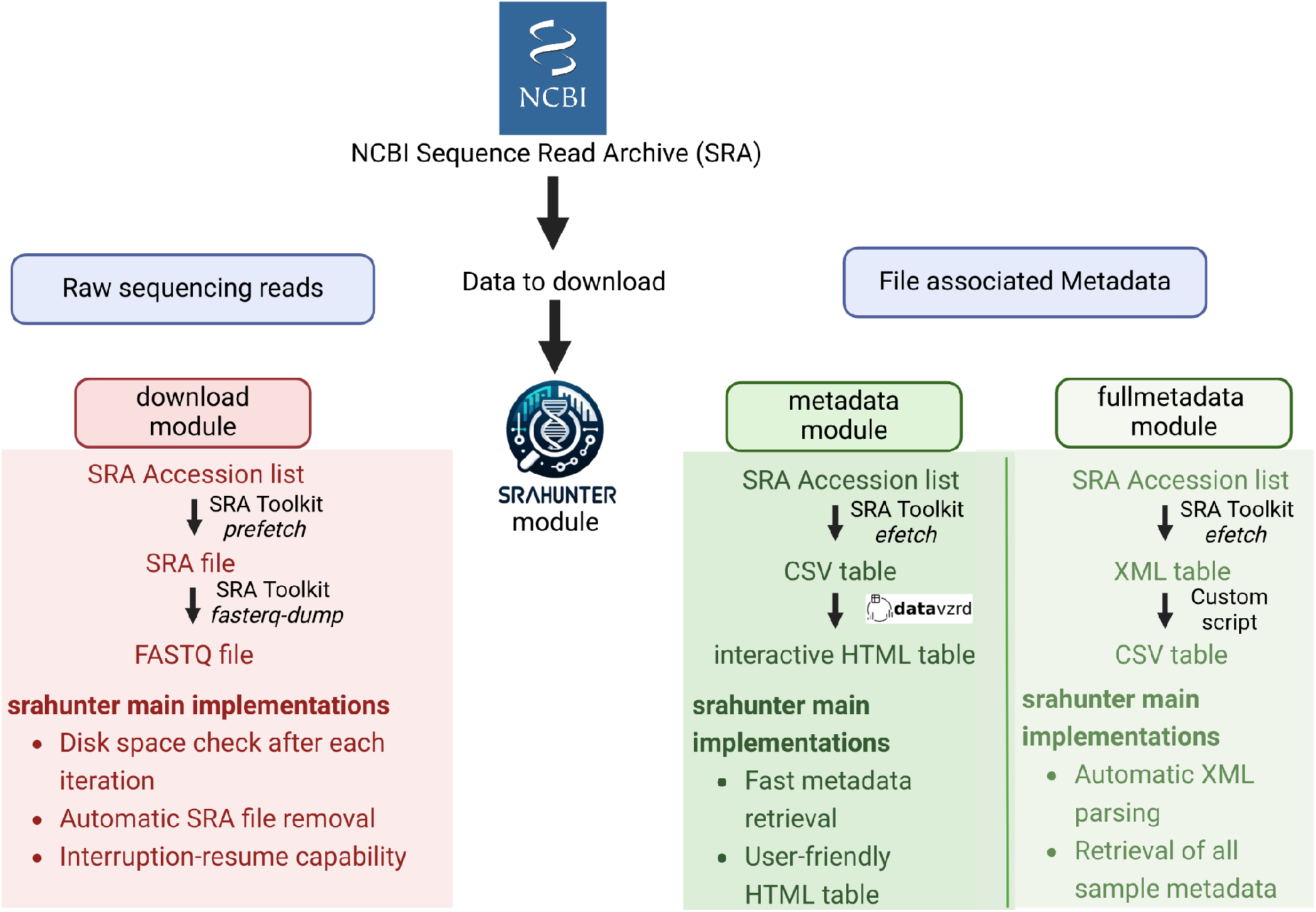
The srahunter workflow. The picture depicts the work steps and main implementations for each of the three different modules.

The download module automatically removes the files in the native SRA format after successful conversion into FASTQ files, data caching, download resuming, and error reporting in case of failed downloads (failed_list.csv file). Users can provide input parameters, such as number of threads (default: 6), download directory (default: tmp_srahunter folder in the current directory), final output path (default: current directory), and maximum SRA file size (default 50 GB).

The metadata module features fast metadata retrieval, integrating the *Entrez direct* functionality and customizable output in CSV and HTML format. The tool supports an input list containing Run accession IDs, whereas Bioproject accessions are not yet supported.

The fullmetadata module allows the user to get all the associated metadata from a list of Run accession numbers, exploiting the Entrez direct functionality, and parsing the XML output. In this case, the final output is represented by a table in CSV format.

### 2.1. The HTML table improves the output readability

In the case of the metadata module as default output, in addition to the CSV file, an interactive HTML table will be produced. This interactive table is produced with the *datavzrd* tool (https://github.com/datavzrd/datavzrd), exploiting different features to facilitate the understanding of the downloaded metadata (Figure 2a). For each column, by clicking on the magnifying glass, it is possible to filter the data, whereas by clicking on the depicted chart, it is possible to produce a chart that summarises the distribution of the column data, and this feature is available for both categorical and numeric variables (Figure 2b and Figure 2c).

**Figure 2.**
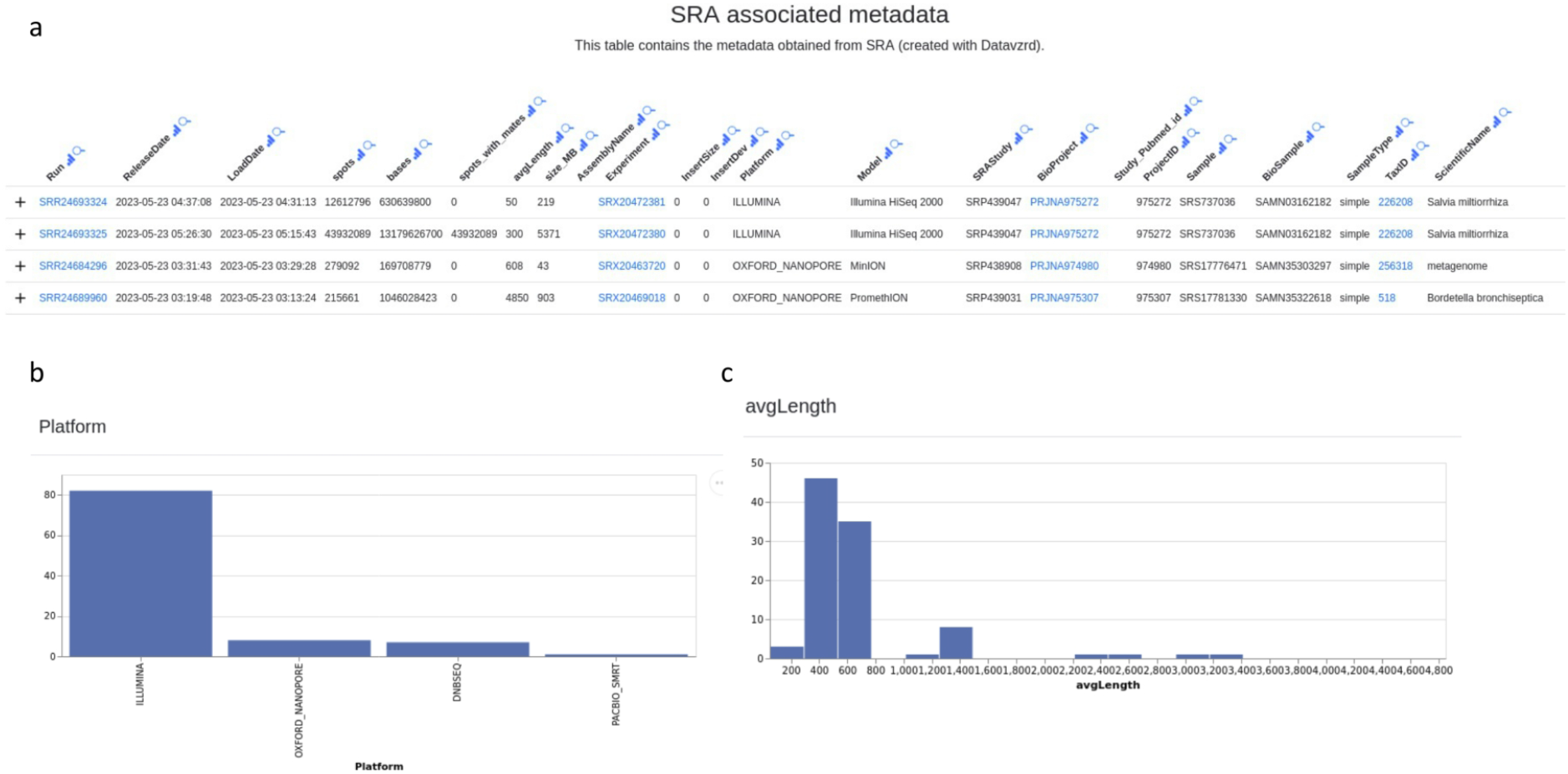
HTML output provided by srahunter. a) An example of the srahunter HTML table is provided: close to each column header a magnifying glass and a schematic chart allow to filter and produce a bar plot of the data distribution, respectively. The bar plot can be produced for both categorical variables (e.g., platform used to sequence the sample in panel b) and numeric variables (e.g., average length of the produced reads in panel c). The list of SRA accession IDs is available in Supplementary file 1.

### 2.2. Comparison with existing tools

For the metadata and full metadata modules, we compared the speed of metadata retrieving with two popular tools, namely *Kingfisher* (https://github.com/wwood/kingfisher-download) and *ffq* [6]. Runtime depends on software, environment, connection speed, and input (machine characteristics are reported in the Supplementary file 2). The *ffq, Kingfisher*, and *srahunter* tools were compared by recording the time needed for retrieving metadata of 1, 10, 100, and 1000 samples using the bash time function (The list of SRA accession IDs is available in Supplementary file 1). The results show that *Kingfisher* and *srahunter* have a significantly lower data retrieval time than ffq (Figure 3a), with no significant differences between the *srahunter* metadata and fullmetadata modules. To better highlight the possible differences between the three tested tools, we compared the median time to compute one sample, which is comparable among *srahunter* metadata, *srahunter* fullmetadata, and *Kingfisher* with a value of 0.13s, 0.17s, and 0.12s respectively, whereas in the case of ffq module the time per sample is slightly higher (0.98s) (Figure 3b).

**Figure 3.**
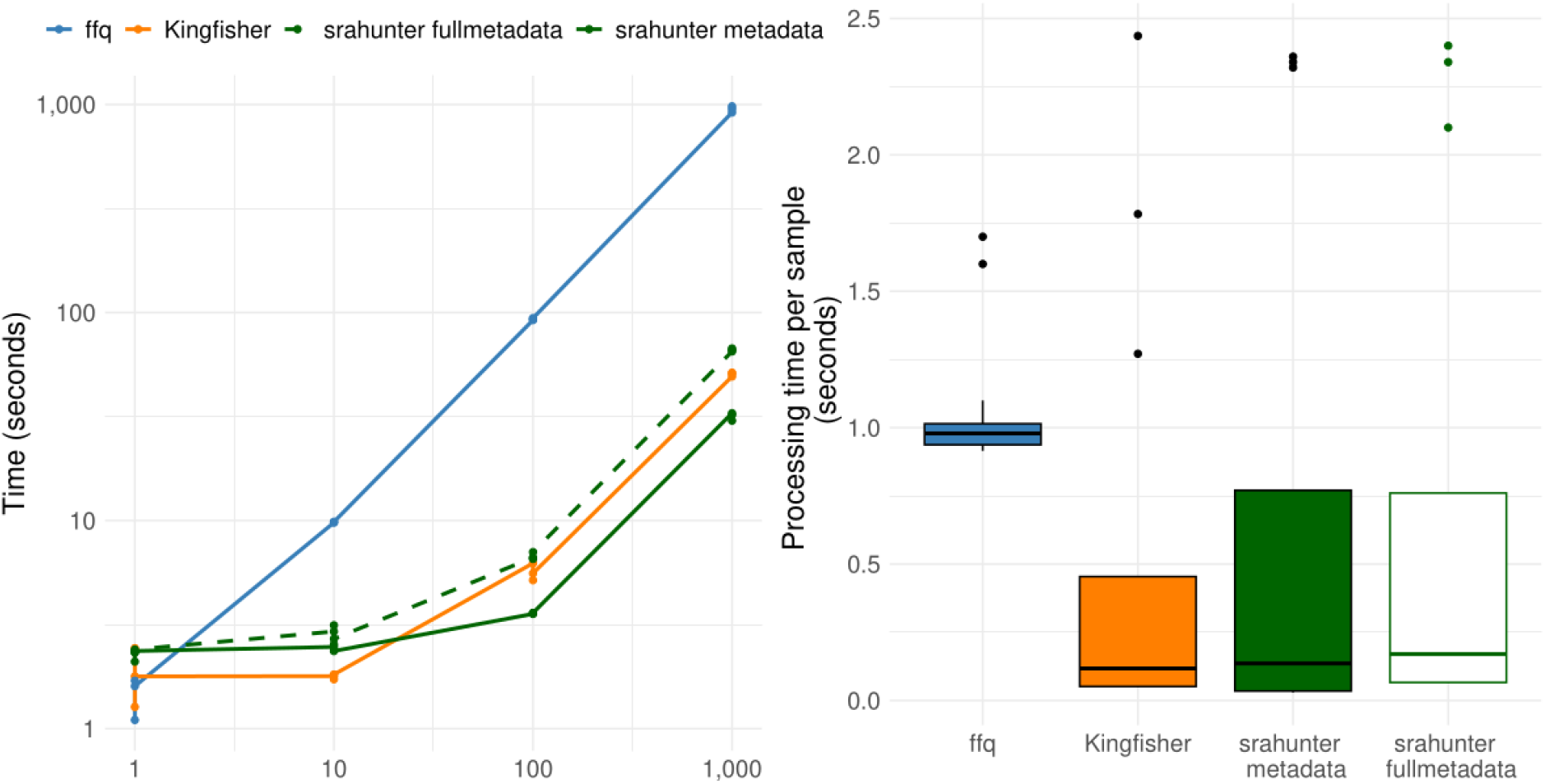
Performance of three different tools in the retrieval of SRA metadata. In the figure panels, ffq, Kingfisher, and srahunter are depicted in blue, orange, and green, respectively. a) The line plot indicates the average time needed to obtain SRA metadata of 1, 10, 100, and 1000 samples with ffq, Kingfisher, and srahunter. The different modules of srahunter, i.e., metadata and fullmetadata, are reported as an unbroken and dotted line, respectively. b) The boxplot shows the average time (seconds) required to analyse one sample. The different modules of srahunter, metadata and fullmetadata are reported as a green filled and a white filled boxplot, respectively.

### 2.3. Example usage

As part of the bioconda repository, the installation of srahunter can be done using the typical command of conda. *srahunter* is well integrated with NCBI SRA and, as an example of its application, we downloaded data and metadata using the default SRA Accession list. The list of commands required for the installation and for exploring the different srahunter functionalities are:

*conda install -c bioconda srahunter*

*srahunter metadata -i SraAccList*.*csv*

*srahunter fullmetadata -i SraAccList*.*csv*

*srahunter download -i -t 12 SraAccList*.*csv*

Of note, *srahunter* is designed to be as simple as possible. That is why for most applications the -i flag can be enough. In the case of the download module, we strongly recommend using the -t flag, which allows a parallel processing of the input files. All the different flags for the three modules are reported on the dedicated Github page.

### 2.4. Scalability and Integration with Other Pipelines

One of the standout features of *srahunter* is its scalability. As the size and complexity of SRA continue to grow, the ability of *srahunter* to manage large datasets efficiently becomes increasingly important. Whether researchers are working with small pilot studies or large-scale sequencing projects, *srahunter* offers the flexibility needed to scale operations without compromising on speed or usability. Additionally, its modular design allows for the integration into existing bioinformatics pipelines, which enhances its utility in workflows involving downstream analyses.

### 2.5. Implementation

*srahunter* is written in Python, with the core functionalities based on *Sra toolkit* and *Entrez direct. srahunter* includes three main modules: download, metadata, and fullmetadata. The download module is designed to retrieve sequencing data from SRA in FASTQ format. The metadata module allows the download of run-associated metadata from SRA and provides, using *datavzrd*, an interactive HTML output. The full metadata module enables the download of the complete set of run-associated metadata. The conversion of SRA to FASTQ files can be parallelized across multiple cpu-cores. Storage availability is constantly checked during downloads, and all processes can be visually monitored through terminal messages and progress bars.

## 3. Conclusion

The ability to efficiently manage large sequence datasets will be critical for future research, such as population-scale genomic studies and, in general, extensive analyses of DNA and RNA datasets. The user-friendly features of *srahunter* and its efficiency in handling complex metadata can improve the reproducibility and transparency of HTS-based studies, both essential for ensuring the reliability of scientific findings. Looking ahead, the functionalities of *srahunter* can be expanded, for instance, the incorporation of additional accession types, such as Bioprojects, would allow researchers to retrieve data across broader organizational levels, further streamlining dataset acquisition.

## Code availability

*srahunter* repository is available on Github https://github.com/GitEnricoNeko/srahunter installation of srahunter requires Conda. More details can be found in the repository.

## Acknowledgments

We thank Nicolò Fogal and Dr. Gaetan Thilliez for tool beta testing. We would like to thank the HPC infrastructure CAPRI (Calcolo ad Alte Prestazioni per la Ricerca e l’Innovazione, University of Padova Strategic Research Infrastructure Grant 2017) for providing the computational power needed for tool testing.

