## Supplementary file 2 for "*srahunter*: a user-friendly tool to speed up and simplify data downloading from NCBI SRA"

**Supplementary file 2.** Technical specification of the machine used for the tool comparison

**CPU:** 13th Gen Intel(R) Core(TM) i9-13900KF

**GPU**: NVIDIA RTX A4000

**SSD:** MZ-77Q4T0BW SAMSUNG 870 QVO SATA 2.5" SSD 4TB

**RAM:** 4X Transcend JM3200HLE-32G 32GB DDR4 3200MHz

**Mother Board:** Gigabyte B660 DS3H DDR4 Intel B660 LGA 1700 ATX

**Download:** 583.53 Mbit/s

**Upload:** 757.44 Mbit/s
